# Clinical implementation of single-cell RNA sequencing using liver fine needle aspirate tissue sampling and centralized processing captures compartment specific immuno-diversity

**DOI:** 10.1101/2021.11.30.470634

**Authors:** Alex S. Genshaft, Sonu Subudhi, Arlin Keo, Juan D. Sanchez Vasquez, Nádia Conceição-Neto, Deeqa Mahamed, Lauke L. Boeijen, Nadia Alatrakchi, Chris Oetheimer, Mike Vilme, Riley Drake, Ira Fleming, Nancy Tran, Constantine Tzouanas, Jasmin Joseph-Chazan, Martin Arreola Villanueva, Harmen J. G. van de Werken, Gertine W. van Oord, Zwier M.A. Groothuismink, Boris J. Beudeker, Zgjim Osmani, Shirin Nkongolo, Aman Mehrotra, Jordan Feld, Raymond T. Chung, Robert J. de Knegt, Harry L. A. Janssen, Jeroen Aerssens, Jacques Bollekens, Nir Hacohen, Georg M. Lauer, Andre Boonstra, Alex K. Shalek, Adam Gehring

**Author notes:** Corresponding Author Adam Gehring Toronto Centre for Liver Disease PMCRT 10-356 101 College St Toronto, ON, M5G1L7 Canada. Denotes shared first authorship (Sonu Subudhi, Arlin Keo, Juan D. Sanchez Vasquez, Nádia Conceição-Neto). Denotes shared senior authorship (Georg M. Lauer, Andre Boonstra, Alex K. Shalek, Adam Gehring).

## Abstract

Blood samples are frequently collected in human studies of the immune system but poorly represent tissue-resident immunity. Understanding the immunopathogenesis of tissue-restricted diseases, such as chronic hepatitis B, necessitates direct investigation of local immune responses. We developed a workflow that enables frequent, minimally invasive collection of liver fine-needle aspirates in multi-site international studies and centralized single-cell RNA sequencing data generation using the Seq-Well S^3^ picowell-based technology. All immunological cell types were captured, including liver macrophages, and showed distinct compartmentalization and transcriptional profiles, providing a systematic assessment of the capabilities and limitations of peripheral blood samples when investigating tissue-restricted diseases. The ability to electively sample the liver of chronic viral hepatitis patients and generate high-resolution data will enable multi-site clinical studies to power fundamental and therapeutic discovery.

---

Understanding the immunopathogenesis of tissue-restricted diseases, and monitoring tissue specific effects of treatment, necessitates direct investigation of local immune responses. However, the only type of human research material appropriate for this purpose that can be collected regularly, and electively, is blood, which often does not mirror tissue-resident immunity. Tissue access for research is limited by ethical and practical considerations, creating a significant obstacle for exploratory human studies. Even when elective tissue specimens can be obtained, clinical sites may lack the personnel and technical infrastructure required to process and handle the material onsite for the optimal generation of state-of-the-art scientific data. Therefore, procedures that enable elective sampling of tissues and generation of high-resolution data at multiple clinical sites would empower both fundamental and therapeutic discoveries.

Chronic liver diseases pose a major public health threat with over 800 million people worldwide at risk for liver cirrhosis and cancer, including over 290 million chronically infected with Hepatitis B virus (HBV)^1, 2^. Chronic hepatitis B (CHB) is a highly heterogeneous disease with different phases characterized by variable viral loads and liver inflammation^3^. Available antiviral therapies effectively limit viral replication but rarely lead to self-sustained functional cure^4^. Currently, there are no known peripheral blood biomarkers to monitor the anti-HBV response in the liver to predict disease progression or functional cure^5^. This is a significant knowledge gap that could be addressed through elective liver tissue sampling and next generation single-cell profiling methods, such as single-cell RNA sequencing (scRNAseq).

Liver tissue for research is typically available through surgical resections or percutaneous needle biopsies. However, these procedures are strictly dependent on clinical need and thus cannot provide the frequent, elective sampling necessary to understand the immune mechanisms at play in a dynamic disease like chronic hepatitis B. Fine-needle aspirates (FNA) of the liver are collected using a 25 gauge (G; 0.51mm outer diameter) needle, which is significantly smaller than needles used for standard liver biopsies (16-18G; 1.65–1.27mm outer diameter) and even standard venipuncture (21G; 0.82mm outer diameter). As a result, the FNA technique poses minimal risk and discomfort to participants. The improved safety profile of liver FNAs enables elective liver sampling for research purposes, including longitudinal clinical studies with less than 2 weeks between samplings^6, 7^. However, compared to more invasive needle biopsies, liver FNAs yield fewer cells (<100,000 cells)^8, 9^ that are best used immediately onsite in assays such as scRNAseq to provide the most accurate representation of the liver immune environment. However, variations in specimen handling and downstream processing, before and during scRNAseq library generation and sequencing, can introduce batch effects that could mask important biological variation after integration of multi-site generated data.

To capitalize on the sampling opportunities enabled by FNAs and address the obstacles of site-specific variability, we developed and optimized a workflow that enables international, multi-site collection of liver FNAs and centralized scRNAseq library generation. First, we developed metrics to assess FNA quality, followed by comparison of two distinct methods for performing scRNAseq on the collected cells, the Seq-Well S^3^ and 10x Genomics 3’ v2 single cell platforms. Both technologies performed similarly in key technical metrics but the cassette-based picowell array of the Seq-Well S^3^ platform did not require advanced technology onsite and captured cell populations lost in the 10x Genomics platform, such as granulocytes. To minimize the impact of processing at different sites, we further optimized a method to freeze and ship loaded arrays, simplifying onsite processing for clinical workflows and facilitating enhanced reproducibility through centralized whole transcriptome amplification, library preparation, and sequencing. Finally, we rigorously analyzed matched blood and liver FNA samples collected at four international sites and highlighted the impact of the liver microenvironment on T cells and our ability to capture liver macrophages with the FNA procedure. This optimized workflow, which can be deployed in multi-site international studies, establishes a precedent for other FNA-accessible tissues to investigate tissue-resident immune responses at the single-cell level without the need for sophisticated onsite technical infrastructure.

## Results

### Fine needle aspirates to assess the diversity of intrahepatic immune cells

Unlike core biopsies that cut a tissue cylinder, FNA sampling uses simultaneous negative pressure and forward motion of the syringe to aspirate cells. This introduces the potential for significant blood contamination if the needle engages hepatic blood vessels. Because of the collection method, the cellular composition of liver FNAs lies between peripheral whole blood and core biopsies^10^. To ensure consistent sampling of intrahepatic tissue, it is important to establish defined procedures that minimize blood contamination and to objectively assess the extent of contamination for each sample.

To this end, we established a standardized protocol for obtaining FNA material with the least possible blood contamination, followed by immediate assessment of sample quality. The protocol prescribes four individual FNA passes, each of which typically displayed a different degree of peripheral blood contamination based on visual inspection (Fig 1a). To measure the degree of peripheral cells contained within a specimen, one can quantify the presence of naïve T cells by flow cytometry, which are typically excluded from solid tissues^8^. However, flow cytometry requires significant time to prepare, a large fraction of the collected sample, and infrastructure readily available at different sites within a clinical trial. Therefore, we tested whether an optical density (OD) test to measure red blood cell (RBC) contamination would deliver comparable results while minimizing sample use and processing time. We compared OD450 measurements with conventional immunological profiling using flow cytometry on different FNA passes from individual participants (Suppl Fig 1). As the OD450 value increased, the CD4:CD8 ratio inverted. CD8 T cells dominated in samples with a low OD450 (typically observed in tissues) to a predominance of CD4 T cells in high OD450 samples, as is characteristic for the blood (Fig 1b)^6^. The value of the OD450 measurement was further validated by finding a significant positive correlation between OD450 and the frequency of both naïve CD4 and CD8 T cells (Fig 1c,d). Furthermore, there was a significant negative correlation between OD450 measurements and frequencies of mucosal-associated invariant T (MAIT) cells (Fig 1e), which are known to be compartmentalized to the liver^11^. These data demonstrate that RBC content measured by OD450 is a robust indicator of peripheral blood contamination in FNA passes and supports the use of this simple, quantitative, sample-sparing test, to select FNA samples with predominantly liver-derived cells for use in subsequent assays.

**Figure 1.**
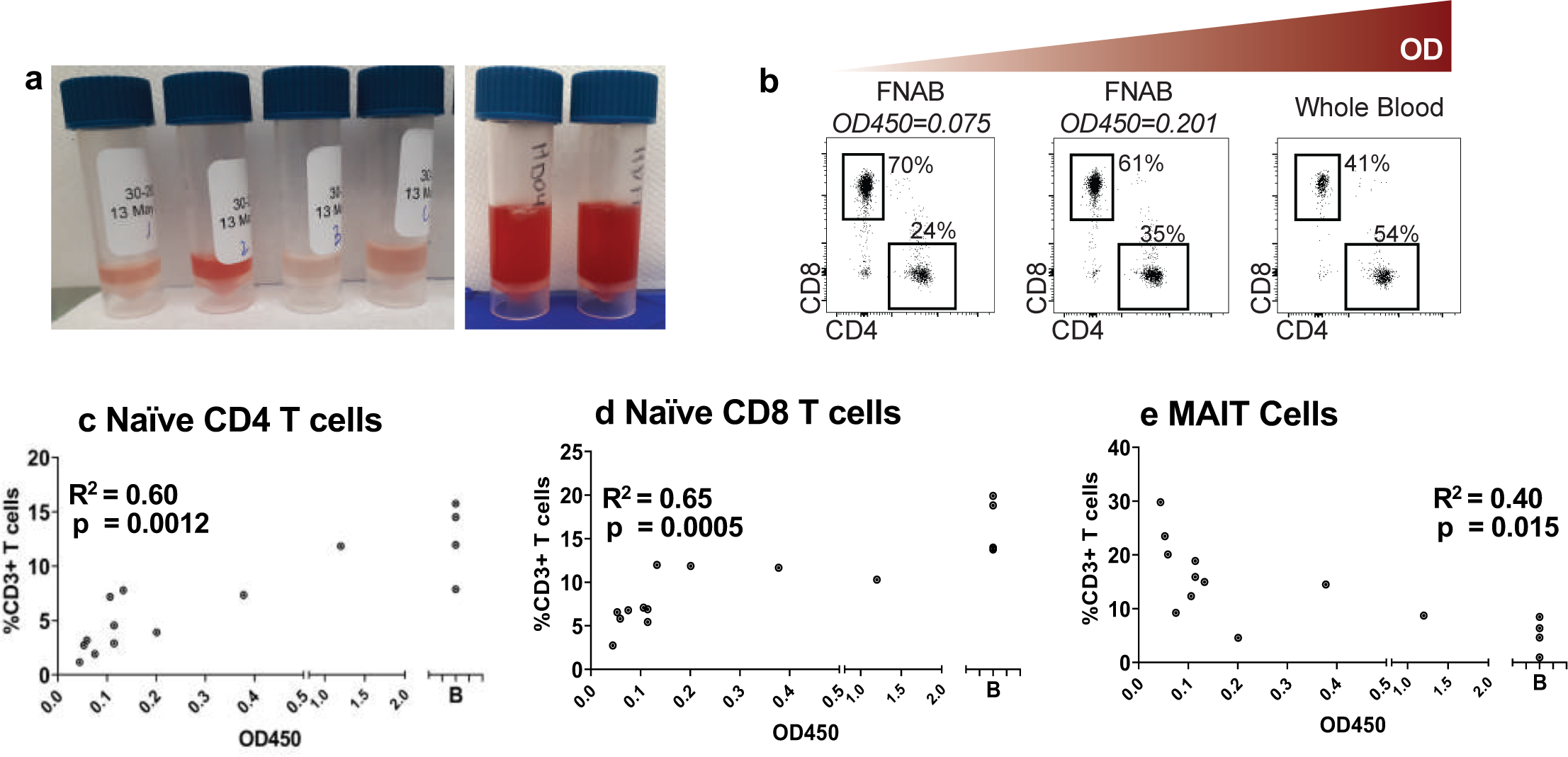
Quantification of RBC contamination of liver FNAs. **(a)** Images showing variable RBC contents of individual FNA passes. **(b)** CD4:CD8 ratio in FNA passes from single patient with increasing OD450 values. Frequency of **(c)** naïve CD4 T cells **(d)** naïve CD8 T cells and **(e)** MAIT cells compared to OD450 values of each FNA pass. Analysis done using Pearson pairwise correlation.

### Comparing scRNAseq technology platforms for analysis of the cellular and transcriptional profile of human liver FNAs

Having optimized the approach for collecting and assessing intrahepatic tissue samples via FNA, we next sought to identify the best means of comprehensively profiling the FNA samples to investigate total cellular diversity. The recent advent of high throughput single-cell RNA sequencing technologies enables genome wide transcriptomic profiling of single cells within complex cellular mixtures to characterize the molecular profiles and intracellular circuits that define them. Both reverse-emulsion droplet and picowell-based high throughput scRNAseq platforms have been introduced that are amenable to low input samples. On one hand, the 10x Genomics Chromium system, like the Drop-seq and inDrops platform, uses a reverse emulsion microfluidic system to co-capture uniquely barcoded oligo-dT beads and cells and performs the initial steps of cell lysis and mRNA capture for library generation onsite^12–14^. On the other hand, the picowell based platforms, such as Seq-Well S^3,15^, and more recently BD Rhapsody^16^, use picowells to isolate individual cells in place of reverse emulsion to perform lysis and mRNA capture for library generation. Both the Seq-Well S^3^ and 10x Genomics platforms have been successfully used to analyze peripheral blood, and to some extent, digested tissue^17–20^. However, the degree of cell capture, cell type representation, sequencing depth and quality, and the robustness of data generation across multiple sites from primary human liver tissue samples is not known.

To benchmark the two scRNAseq platforms, four FNA passes were collected from each of four volunteers. The highest quality passes were pooled and then analyzed in parallel using Seq-Well S^3^ and the 10x Genomics 3’ version 2 (10x v2) platforms. After filtering for low-quality cells based on a minimum 300 genes and 500 UMIs per cell, the number of transcripts (p=0.044), number of genes captured per cell (p=0.013) and cell capture from 15,000 cell input (p=0.039) was significantly higher in liver FNAs using Seq-Well S^3^ (Suppl Fig 2a-c). For the peripheral blood, only the number of transcript (p=0.039) showed a significant difference between Seq-Well S^3^ and 10x Genomics (Suppl Fig 2d-f).

UMAP clustering of the Seq-Well S^3^ and 10x 3’ v2 datasets readily identified major lymphocyte (T cells, B cells, NK cells) and myeloid (monocytes, macrophages, dendritic cells (DC)) cell types using lineage-specific markers (Fig 2a-d). Using identical samples loaded onto the Seq-Well S^3^ and the 10x 3’ v2 system, we compared cell type frequencies measured by the two platforms by calculating the number of each cell type divided by the total cell count of sequenced cells passing quality thresholds (Fig 2e). In this head-to-head comparison, 10x v2 captured significantly more γδ T cells (p = 0.017) whereas Seq-Well S^3^ captured significantly more granulocytes (p=0.008) and was capable of efficiently capturing both blood and liver neutrophils, which were undetectable in the 10x 3’ v2 dataset (Fig 2e). In addition to neutrophils, a cluster of regulatory T cells (*CD3*, *CTLA4*, *IL2RB*, *FoxP3*) was unique to the Seq-Well S^3^ dataset (p=0.0009), when compared to data obtained using the 10x 3’ v2 kit. Very few high-quality non-immune cells, such as hepatocytes, were captured using either method. This is likely intrinsic to the liver FNA collection method since the size of the FNA needle may not effectively mobilize hepatocytes and the negative pressure of aspiration could cause physical stress on the fragile cells.

**Figure 2.**
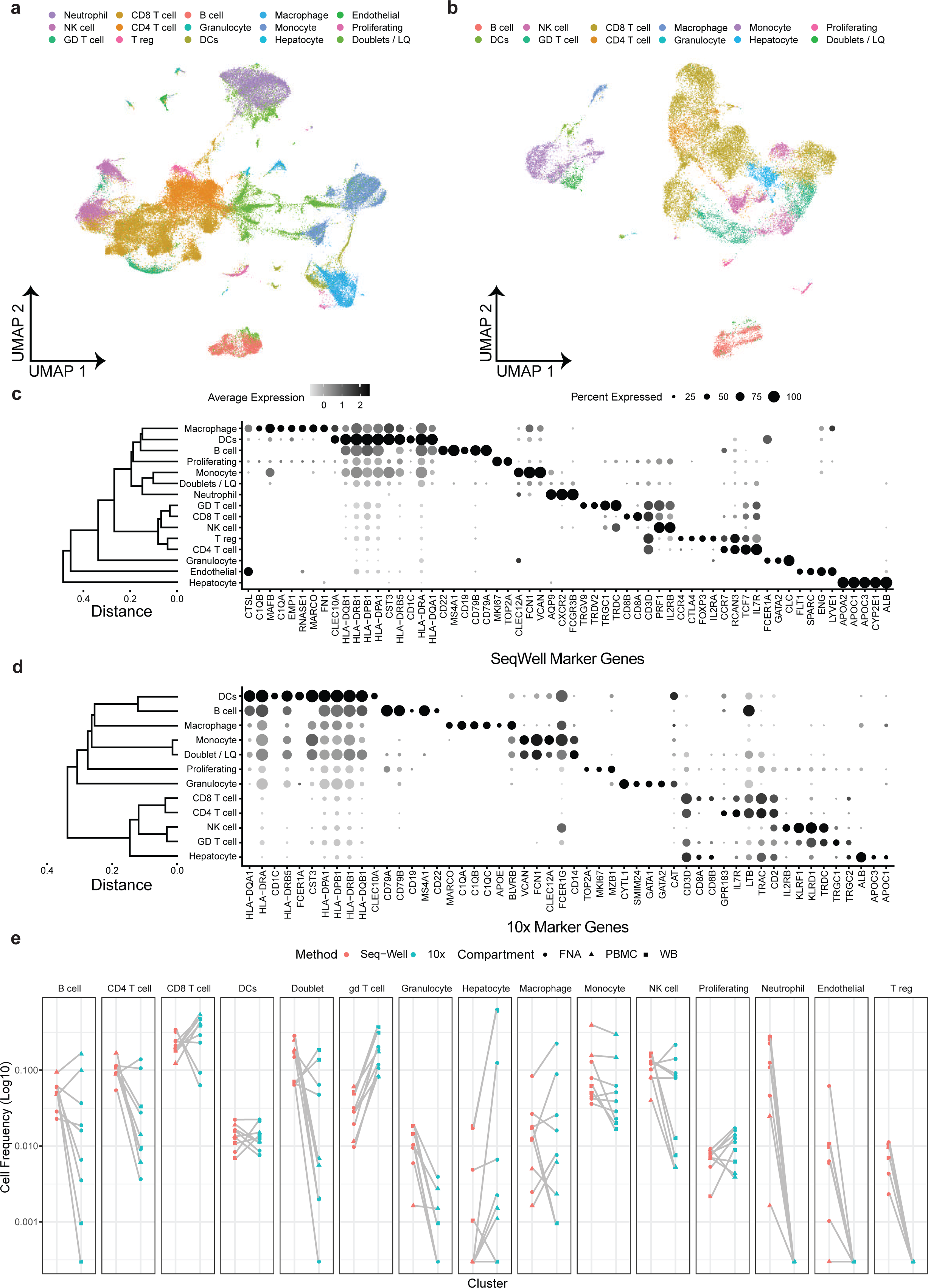
10x 3’v2 vs Seq-Well S^3^ comparison. UMAPs of all data from **(a)** Seq-Well (n = 67,947) and **(b)** 10x cell types (n = 25,473). Dot plot of cell type marker genes for **(c)** Seq-Well and **(d)** 10x. **(e)** Cell type frequencies compared across matched samples (gray lines) for Seq-Well S^3^ (red) and 10x (blue). Samples from FNA, PBMC, and WB are circles, triangles and squares, respectively.

### Cryopreservation of Seq-Well S^3^ arrays for centralized sequencing library generation

Based on the representative cell types captured, and that the picowell-based Seq-Well S^3^ system required minimal specialized equipment onsite, we selected this approach for further evaluation. To enable work with complex clinical samples, we tested a protocol that would allow for centralized processing of critical steps. Specifically, we piloted an approach to freeze Seq-Well S^3^ arrays immediately after cell loading and membrane sealing. If successful, this would enable loading of freshly isolated cells on picowell arrays and perform all subsequent steps – including lysis, hybridization, reverse transcription, exonuclease digestion, whole transcriptome amplification, library preparation and sequencing – centrally to minimize onsite processing and potential batch effects.

To test the feasibility of this approach, we loaded FNA and PBMC samples from six participants from 3 different sites onto parallel Seq-Well S^3^ arrays up to membrane sealing. One array was processed directly on site up to the overnight reverse transcription step and the other one was frozen at -80 °C and shipped on dry ice to the central processing lab. Freezing and shipping after cell loading yielded equivalent number of transcripts, genes per cell and total cell recovery compared to processing onsite in both the liver FNA (Fig 3a-c) and peripheral blood samples (Fig 3d-f). Equally important, fresh and frozen array datasets clustered together seamlessly, without data integration, with only a maximum of seven consensus differentially expressed genes (consensus = expressed in >50% of comparisons) observed in CD8 T cells between fresh and frozen arrays (Fig 3g). Similarly, arrays from individual patients showed similar cellular diversity when processed via either protocol (Fig 3h). The ability to perform library generation and sequencing at a central processing lab not only increases the robustness of the assay, but also enables clinical studies in low resource settings without specialized equipment such as thermocyclers 10x Chromium controllers.

**Figure 3.**
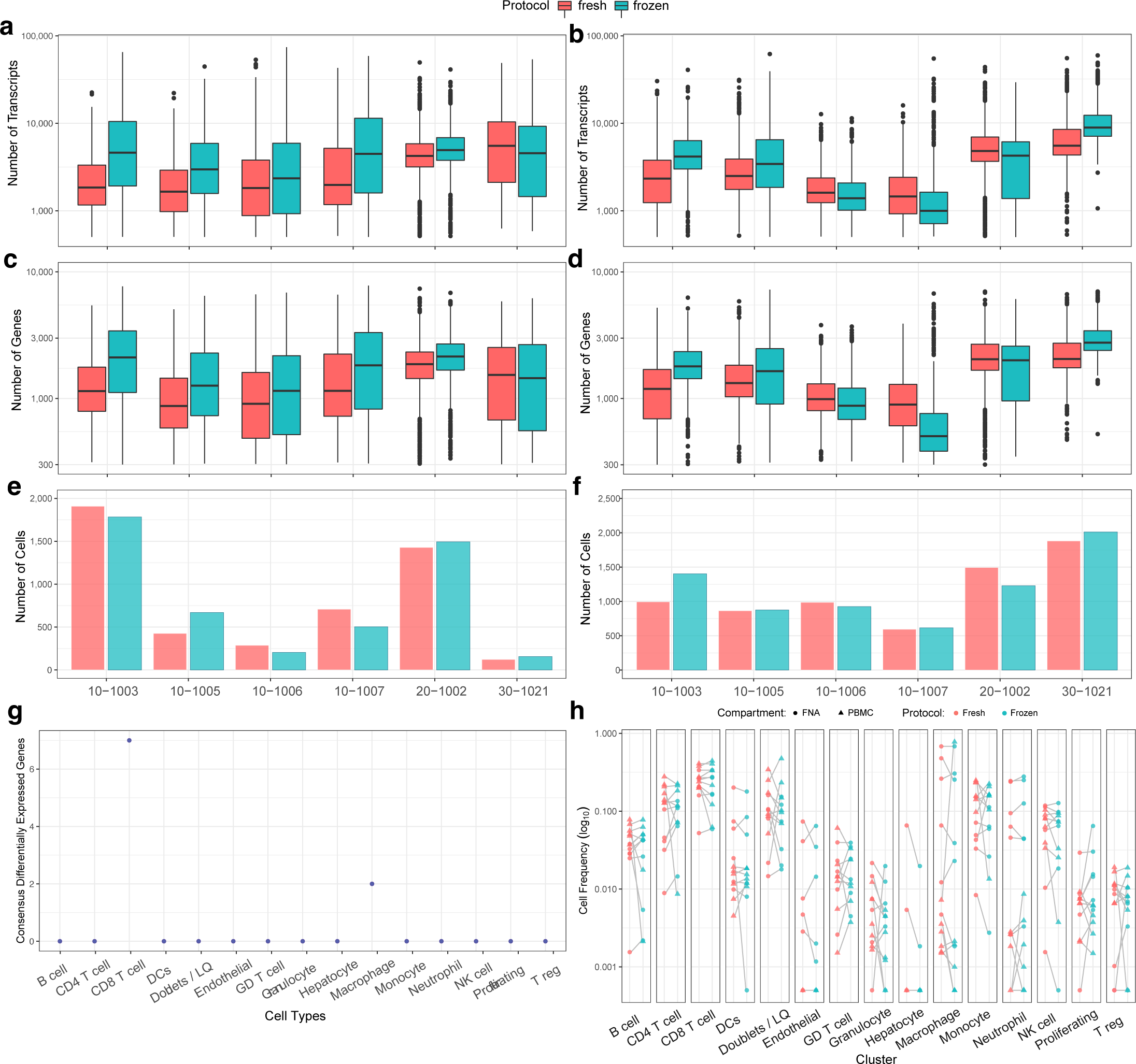
Data comparison of freshly processed vs. frozen arrays. **(a)** number of transcripts/cell **(b)** number of genes/cell and **(c)** number of cells captured in liver FNAs from six different patients. **(d)** number of transcripts/cell **(e)** number of genes/cell and **(f)** number of cells captured in peripheral blood from six different patients. **(g)** Number of differentially expressed genes in matched cell types between freshly processed and frozen arrays. **(h)** Comparison of cell type capture between freshly processed and frozen arrays in matched samples.

### Implementing the optimized FNA processing workflow to examine liver-resident immune cell diversity

To broaden the dataset for a more robust appreciation of the cellular subsets and features recovered using the FNA procedure, we used the optimized workflow to characterize liver FNA samples from 13 CHB participants with inactive disease and 3 healthy participants. Blood and FNA samples from all participants were subjected to scRNAseq with the Seq-Well S^3^ platform. After preprocessing, we recovered a total of 66,446 high-quality cells, which were subclustered and analyzed in a lineage-specific manner to provide a comprehensive, comparative map of the intrahepatic immune landscape and to test whether transcriptional differences are apparent between immune cells from the blood and liver.

### Subset enrichment and transcriptional adaption of CD8 T cells in the liver

T cells are a major focus of interest in HBV infection given their critical role in viral control and liver disease progression. Previous observations indicate that the composition and phenotypic profiles of liver T cells is significantly distinct from what is found in the blood^21^. Subclustering data from 16 participants revealed 5 distinct CD8 T cell populations (Fig 4a). The top 10 differentially expressed genes discriminating each cluster are shown in Figure 4b (a full list of gene markers is available in Supplementary Table 1). Analysis of all cluster defining markers revealed that these are enriched for signatures of prototypic CD8 T cell subpopulations known from prior studies of infection and cancer^22^. Cluster 0 (“GZMB”) was characterized by a strong expression of granzymes B (*GZMB*) and H (*GZMH*), together with the chemokine receptor *CX3CR1* and other markers typically found in effector T cells^22, 23^. Cluster 1 (“GZMK”) was dominated by expression of *GZMK* together with the transcription factor *EOMES* and the chemokine receptors *CCR5* and *CXCR6*. This population closely resembles transitional or precursor CD8 T cells previously described in liver cancer^22^. The key feature of cluster 2 (“*NR4A2”*) is the expression of the transcription factor *NR4A2* that has been associated with dysfunctional CD8 T cells^24^. Cluster 3 (“CCR7+ TCF7+”) expressed the chemokine receptor *CCR7* and transcription factor *TCF7,* together with the regulatory protein *LEF1* and the adhesion molecule *SELL*, a combination that is characteristic of both naïve and naïve-like, or stem-like, CD8 T cells^25^. Finally, cluster 4 (“MAIT”) co-expressed classical genes associated with MAIT cells such as the NK receptor *KLRB1* in addition to *SLC4A10*, *DPP4*, and *IL7R*^26^. Interestingly, the MAIT cell cluster expressed many of the GZMK cluster 1 defining genes, in addition to its own characteristic gene expression signature (Fig 4c).

**Figure 4.**
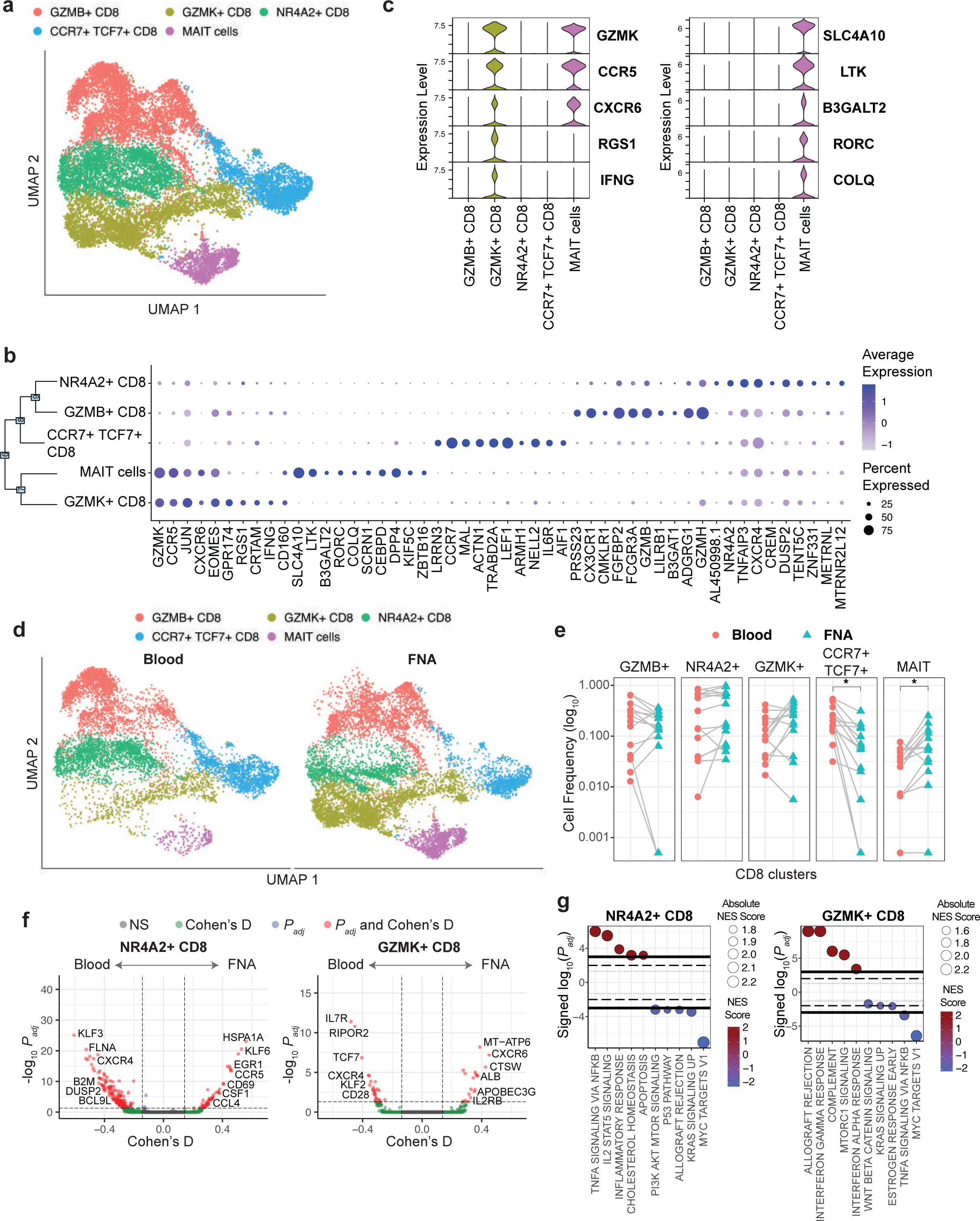
CD8 T cell composition in liver vs blood. **(a)** scRNAseq UMAP for CD8 T cells colored by cluster IDs. **(b)** Dot plot showing top 10 marker genes for each cluster ID. **(c)** Violin plot showing the top 5 marker genes for GZMK+ CD8 and MAIT cells. **(d)** scRNAseq UMAP colored by cluster and split based on tissue of origin, i.e. liver and blood. **(e)** Comparison of cell frequencies between blood and liver within sample (connected through grey lines) for each CD8 cluster. For determining significant differences, Wilcox test with Bonferroni correction was used (adjusted p-value < 0.05). CCR7+ TCF7+ CD8 and MAIT cells are significantly more in blood and liver, respectively. Other clusters are equally distributed across both tissues. **(f)** Volcano plots depicting differences in gene expression between NR4A2+ and GZMK+ CD8 T cells. Positive cohen’s D value suggests higher expression in liver. Cohen’s D cutoff calculated as mean + 2 x standard deviation of cohen’s D values of all genes. **(g)** Hallmark genesets enriched by NR4A2+ and GZMK+ CD8 T cells. X-axis represents signed log10 of adjusted P-value for the genesets, and positive value suggests enrichment in liver.

When comparing the five clusters of CD8 T cells between the blood and FNAs, some clusters were relatively more prevalent in one or the other tissue (Fig 4d). Differences in relative T cell frequencies were apparent for MAIT cells (p=0.039), which are known to be significantly enriched in the liver^11^, confirming that the FNA procedure succeeded in capturing liver-associated immune cells (Fig. 4e). The *GZMK* population was also more prevalent in the liver. In contrast, the naïve/naïve-like *CCR7 TCF7* population was enriched in the blood (p=0.0006), which is expected as naïve cells are mostly excluded from non-lymphatic tissues^27^ (Fig 4e). Deeper comparison of transcriptional profiles between blood and liver within clusters revealed profound differences in gene expression for the *NR4A2* and *GZMK* CD8 T cell clusters (Fig 4f). Enrichment analysis revealed that these differentially expressed genes represent key cellular pathways enriched within liver, including inflammatory response, IL2 and TNF-α signaling in the NR4A2 population whereas IFN-γ and mTORC1 signaling were enriched in the GZMK population (Fig 4g). Overall, despite the potential for blood contamination, liver FNAs captured the liver enriched specific CD8 T cell clusters. The data also demonstrate that tissue residency is associated with distinct transcriptional programs in CD8 T cell subpopulations found in both the liver and blood. This supports the notion that analysis of CD8 T cells from the site of infection is important to fully understand T cell compartmentalization and adaptation.

In contrast to the CD8 T cells, the composition of the CD4 T cell population was less complex, with only 2 distinct clusters (excluding Tregs cluster in Fig 2c). Moreover, further analysis revealed only marginal differences with regards to the relative size of clusters or their transcriptional landscape (Suppl Fig 3).

### Definition of a complex neutrophil compartment in the blood and liver

Animal studies suggest that neutrophils are recruited first in the inflammatory cascade and express matrix degrading enzymes that facilitate immune cell infiltration into the liver parenchyma^28, 29^. There is currently little information on the possible role of neutrophils in HBV pathogenesis as they have been understudied in patients because they are lost upon PBMC isolation and cryopreservation. Once we realized the picowell-based scRNAseq approach efficiently captured neutrophils, we changed our RBC depletion method from density gradient isolation to magnetic bead-based depletion to preserve the neutrophil population. As a result, 7 participants with matched whole blood and FNA were present in the analysis, with the remaining 9 having only PBMCs isolated by density gradient centrifugation. Subclustering of neutrophils yielded six subpopulations (Fig 5a). The loss of neutrophils was apparent in the PBMC samples (Fig 5b). Neutrophils shared an overlapping transcriptional profile with monocytes, such as the *S100A8* and *S100A9* transcripts, but were clearly distinguishable from monocytes due to the lack of *VCAN* and *CD68* (not shown). Using the top five genes from each cluster, we found that five clusters expressed known neutrophil markers such as *CXCR2*, *IL8*, and S100A genes^30, 31^. Cluster 0, IL8(hi) SYAP1-neutrophils, and cluster 4, IL8(hi) SYAP1+ neutrophils expressed the highest level of IL8 (*CXCL8*), which is found in activated neutrophils^32^. Expression of SYAP1, a described target of caspases, distinguished the IL-8+ neutrophils but its role in neutrophil regulation remains unknown (Fig 5c). Both cluster 3, (*MME*) and cluster 5 (*MMP8*) neutrophils expressed genes whose products are stored in secretory granules and aid in the process of neutrophil recruitment to tissues and matrix degradation^33, 34^. However, they expressed different secreted molecules such as *MME* and *FCN1* or *MMP8* and *LCN2* in cluster 3 and 5, respectively (Fig 5c). Cluster 1, IFN-stim neutrophils, displayed a profile consistent with activation by type I interferon, with increased expression of *RSAD2*, *IFIT1*, *MX1*, *ISG15* and *OAS3.* Cluster 2, (*SIGLEC10)* neutrophils*, expressed SIGLEC10,* which has been described as an inhibitory receptor of other immune cells in the context of autoimmunity and tumor escape^35–37^ (Fig 5c).

**Figure 5.**
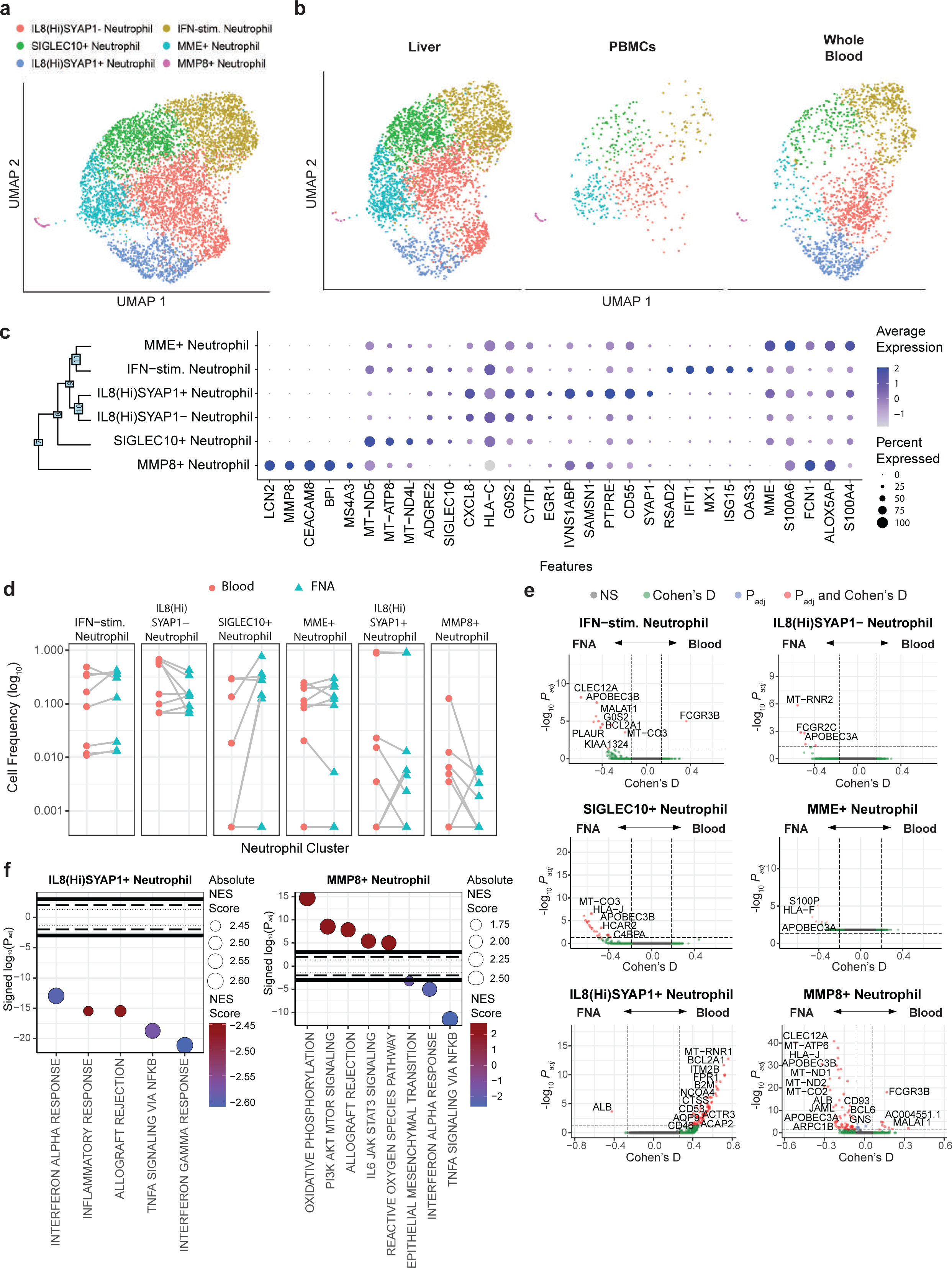
Neutrophil identification. **(a)** scRNAseq UMAP for neutrophils colored by cluster IDs. **(b)** UMAP dimensionality reduction of neutrophils by compartment. **(c)** Dot plot showing top 5 marker genes for each cluster. **(d)** Differential frequency of neutrophil clusters between compartments. Participants with 0 cells within a cell population were excluded. Blood corresponds to whole blood not PBMCs. P-values were Bonferroni corrected for multiple hypothesis testing. **(e)** Volcano plots depicting differences in gene expression between compartments within each cluster. Positive cohen’s D value suggests higher expression in blood. **(f)** Hallmark genesets enriched for clusters with differentially expressed genes between compartments. X-axis represents signed log10 of adjusted P-value for the genesets, and positive value suggests enrichment in liver. IFN-stim. Neutrophil, Interferon stimulated Neutrophil.

Comparing the distribution of neutrophils between the blood and liver for those participants with whole blood samples did not show significant enrichment of any neutrophil cluster in either compartment (Fig 5d). Differential gene expression analysis within neutrophil clusters between liver and blood showed significant genes in four out of the six clusters (Fig 5e). Enrichment analysis of IL8(hi)SYAP1+ neutrophils revealed significant enrichment of pathways associated with type I and II interferons (IFNs) in the blood. Similarly, the pathways that were increased in the blood in *MMP8* neutrophils were also related to type I and II IFNs, whereas pathways elevated in the liver-derived *MMP8* neutrophils were enriched in *IL6* signaling and metabolic processes (Fig 5f). An additional observation was the unique expression of the integrin *CLEC12A* in liver derived neutrophils across multiple clusters (Fig 5e). While their role in the progression of CHB is unclear, the ability to capture neutrophils in the scRNAseq data will allow for deeper investigation and a more in-depth understanding of their role.

### Liver FNAs capture macrophage diversity in the human liver

Macrophages regulate the inflammatory environment in the liver and are associated with HBV-mediated liver inflammation and progression of fibrosis^38–41^. Macrophages exist as a heterogeneous population of embryonically-derived Kupffer cells and monocyte-derived macrophages that lie on a functional spectrum between activating and suppressive^42^. They are tightly bound to the endothelium, and studies that have characterized human macrophages with scRNAseq have used collagenase to digest liver tissue^19^. It was unclear whether FNAs would allow us to capture tightly bound macrophage populations. We noted that macrophage capture using the FNA sampling approach was variable among patients. Therefore, caution should be taken when comparing frequencies of adherent cells, like macrophages, between time points or between patients. Despite the variability in recovery, we identified a clear macrophage cluster within our dataset (Fig 2c). Subclustering of the macrophage population yielded five distinct subpopulations (Fig 6a) that were entirely restricted to the liver (Fig 6b). The macrophage clusters displayed markers previously demonstrated to be shared across macrophage populations including complement components (*C1QA*), *FCGR3A* (CD16), *MARCO*, *CTSS and MSR1* (Fig 6c). However, we identified unique markers capable of distinguishing each population (Fig 6d). Cluster 0, *TIMD4* macrophages, identified by unique expression of *TIMD4*, have been identified as embryonically-derived, or long-lived, tissue macrophages in animal studies and in the human heart and gut^43, 44^. *TIMD4* macrophages highly expressed *LYVE1*, also associated with long-lived tissue resident macrophages^43, 45, 46^, and CD163, a scavenger receptor associated with fibrosis in chronic hepatitis B (Fig 6e)^47^. Cluster 1, *SLC40A1* (ferroportin) macrophages, expressed liver macrophage markers that indicate recent monocyte to macrophage differentiation including *NR1H3* (Liver X receptor alpha, LxRa) and *SPIC* (Fig 6e)^48, 49^. Cluster 2, *C1QA* low macrophages, and cluster 3, *HBB* (hemoglobin) macrophages expressed the lowest level of shared macrophage markers *C1QA* and *MARCO,* and therefore, may represent transient macrophage populations^50–,52^ (Fig 6c). Cluster 2, *C1QA* low macrophages, expressed high levels of monocyte markers *VCAN* and *LYZ*, suggesting these macrophages are in a transitional state of differentiation (Fig 6e)^53^. Cluster 4, HBB+ macrophages, may represent a transient liver macrophage population responsible for clearance of RBCs prior to differentiating to *SLC40A1* macrophages, consistent with the recently differentiated transcriptional profile of *SLC40A1* macrophages^54^. Cluster 4, *CD9* macrophages, expressed *OLR1* and *LGALS3* and were the smallest detectable cluster in the dataset (Fig 6e). *CD9* macrophages represent previously identified scar-associated macrophages found in the cirrhotic liver^20^. These data confirm that the collection of FNAs allows the capture of adherent macrophages and identifies unique macrophage states associated with different stages of liver disease.

**Figure 6.**
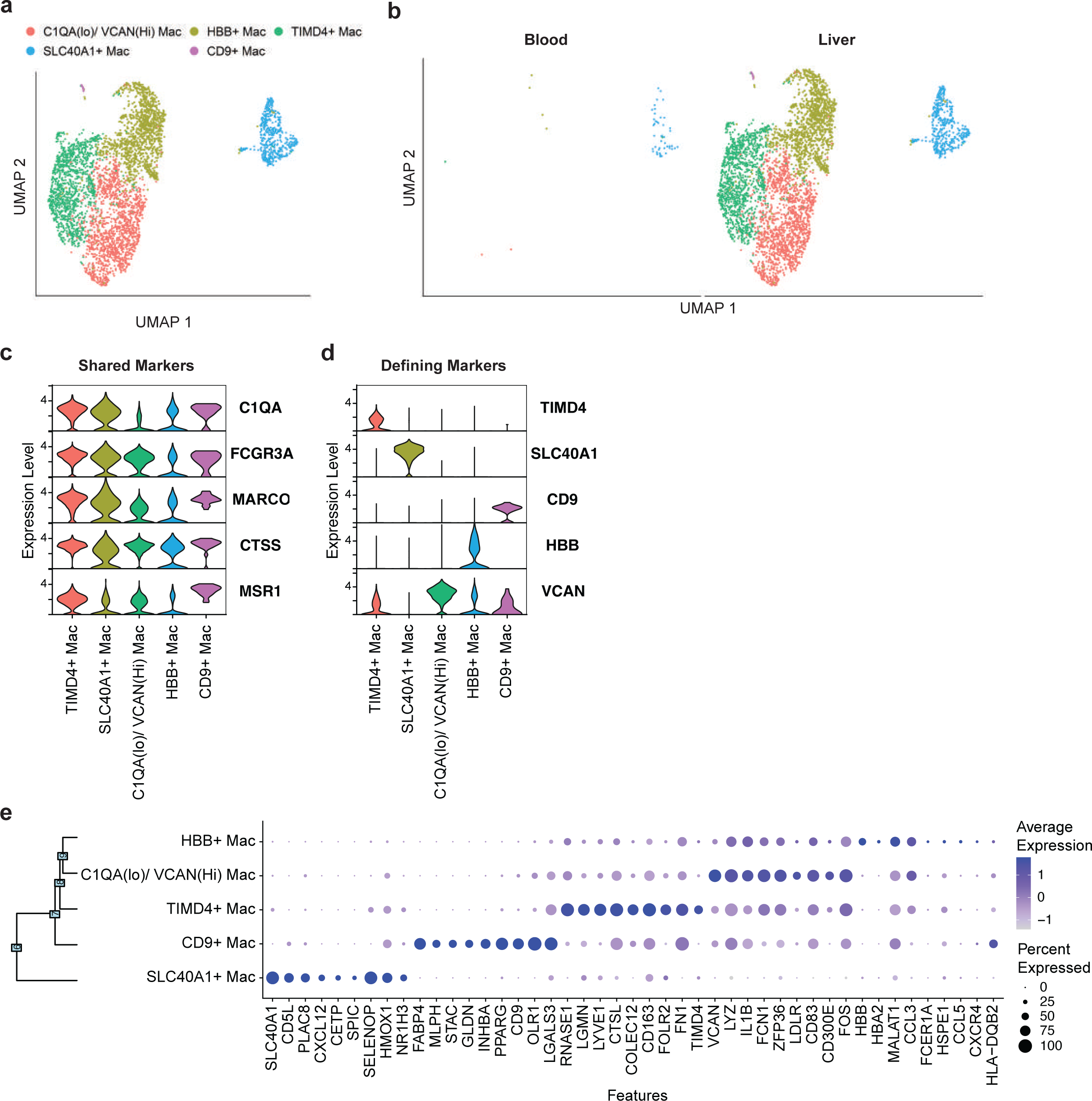
Macrophage identification in liver FNAs. **(a)** scRNAseq UMAP for macrophages colored by cluster IDs. **(b)** UMAP dimensionality reduction of macrophages by compartment. Violin plots of **(c)** macrophage-shared genes and **(d)** unique cluster-defining genes. **(e)** Dot plot showing the top 9 cluster defining genes. All selected genes have an adjusted P value <0.05. Mac, macrophages.

### NK cells display a comparable transcriptional profile between the blood and liver

NK cells are key players in the innate immune response in viral infections, given their ability to inhibit viral replication through cytokines and to recognize and kill virus-infected cells^55^. However, they may also regulate adaptive immune responses both by eliminating virus-specific T cells and through antibody-dependent cell cytotoxicity^56^. NK cells are enriched in the human liver^57, 58^, but the distribution of NK subtypes, and their transcriptional profiles, in the blood and liver of HBV participants has not yet been comprehensively defined. Our dataset consisted of 1,340 peripheral blood NK cells and 3,491 liver NK cells. Clustering analysis of these NK cells revealed two subclusters (Suppl Fig 4a). Cluster 0 expressed low levels of *NCAM1,* which encodes for CD56 (a well-known lineage marker of NK cells), characteristic of CD56-dim NK cells, which have been studied extensively by flow cytometry. Cluster 1 expressed high levels of *NCAM1* representing the CD56-bright NK cells (Suppl Fig 4b). Cluster 0 (“*NCAM1-dim, GNLY*”) were characterized by the cytotoxicity genes GNLY, GZMB, the chemokine receptor CX3CR1 and Fc receptors IIIa (*CD16*). Cluster 1 (“*NCAM1-bright EOMES*”) displayed high expression of the transcription factor *EOMES*, the surface chemokine receptor genes *CXCR6, CCR5*, chemokine ligand *XCL1* and the apoptosis-inducing ligand *TNFSF10* (TNF-related apoptosis inducing ligand (TRAIL; Suppl Fig 4b,c). These transcriptional profiles are consistent with previously reported literature indicating that CD56-dim NK cells display greater cytotoxic activity, while the CD56-bright NK cell subset efficiently produce TRAIL and cytokines/chemokines^56^.

We then analyzed the distribution of each cluster between the blood and liver (Suppl fig 4d). Consistent with prior observations^59^, cluster 1, (“*NCAM1-bright EOMES*”) was significantly enriched in liver FNA compared to the blood while NK cells from Cluster 0 (“*NCAM1-dim GNLY*”) were evenly distributed across the compartments (Suppl Fig 4e). We compared the gene expression for each NK cell cluster between the liver and blood but found no differentially expressed genes in either cluster between the two compartments. Only mitochondrial or non-NK related genes were differentially expressed in *NCAM-bright EOMES* NK cells between the blood and liver (Suppl Fig 4f,g). Therefore, FNA sampling confirmed enrichment of *NCAM-bright EOMES* NK cells in the liver but our data indicates that neither NK cell cluster displayed clear differential expression of NK-related genes between blood and liver.

### Similar B cell composition and transcriptional profiles across the blood and liver

Although B cells have been given less attention than T cells in studies of the immunopathology of HBV, B cells exert some immune control, as demonstrated by HBV reactivation after B cell depleting treatment with anti-CD20 antibodies^60^. Clustering of B cells yielded five subpopulations (Suppl Fig 5a). Cluster 0, *TCL1A* naïve B cells, lacked *CD27* expression and could be distinguished by their expression of *TCL1A*, *FCER2*, *IL4R*, *IGHD*, and *IGHM*. Furthermore, among the naïve subset, we further identified a subgroup of *TCL6* transitional B cells (Cluster 4) by their expression of *PCDH9*, *SOX4*, and *TCL6*^61–63^. Cluster 2, *CD69* naïve B cells, were defined by elevated expression of the NR4A nuclear receptors (i.e. *NR4A1*, *NR4A2*, *and* NR4A3), which have been shown to be rapidly upregulated by B cells upon B cell antigen receptor stimulation^64^, and higher expression of the B cell activation markers *CD83* and *CD69* (Suppl Fig 5b,c)^65, 66^. Within the total memory B cells, Cluster 3 represents *FCRL5*+ atypical memory B cells, which were identified by their expression of *TBX21* (T-bet), *ITGAX* (CD11c), and *FCRL5*^67, 68^. *FCRL5+* atypical memory B cells also overexpressed *SIGLEC6*, which is considered an exhaustion marker for B cells^69^. The remaining population of memory B cells, cluster 1, were defined as *CD27+* classical memory B cells based on their expression of *CD27*, *AHNAK*, and *IGHG1* (Suppl Fig 5b,c)^61^. None of the B cell clusters showed significant enrichment in the blood or liver (Suppl Fig 5d,e) nor did they display significant differences in gene expression within clusters between the liver and blood (Suppl Fig 5f).

### Monocytes display high heterogeneity but not tissue-specific enrichment

A defining feature of monocytes is their plasticity, the ability to differentiate into dendritic cells or macrophages upon exposure to environmental signals. Consistent with their plasticity, we identified a highly diverse monocyte population between the blood and liver, consisting of 10 monocyte subpopulations (Suppl Fig 6a). The 6 most differentially expressed genes discriminating each cluster are shown in Suppl Fig 6b and demonstrate the transcriptional overlap between different monocyte clusters. All monocyte populations expressed *CCR2* and *LYZ* at varying levels. Some clusters showed distinct features. Cluster 4, IFN-stimulated monocytes (IFN-stim cMono), displayed a profile consistent with activation by type I interferon, with increased expression of *IFIT1*, *MX1*, *IFI44L* and *OAS3*. Cluster 8, Intermediate monocytes (Int-Mono), showed elevated MHC-II antigen processing machinery and expression of *FCGR3A* (CD16)^70^. Cluster 9, Non-classical monocytes (ncMono), were identified by expression of *FCGR3A*, *FCGR3B* and chemokine receptor genes *CXCR1* and *CXCR2*^71^. Two classical monocyte populations, Cluster 2 & 3, *CCL3L-* and *CCL3L1+* cMono respectively, displayed a unique transcriptional phenotype characterized by high expression of the cytokines *TNFa* and *IL-1β* (Suppl Fig 6b). These data confirm the known plasticity of monocytes, particularly in disease settings^71^.

Analysis of the distribution of the monocyte subpopulations did not show significant enrichment of any cluster in either blood or liver (Suppl Fig 6c,d). However, two monocyte clusters displayed significant differential gene expression between the compartments: cluster 0, cMono(1), and cluster 1, cMono(2) (Suppl Fig 6e). Enrichment analysis of cMono(1) and cMono(2) showed similar pathways related to a response to cytokine signaling (IL-6, IFN-γ), transcriptional programs (myc), and metabolism (mTor). However, cMono(1) pathways were significantly expressed in the liver while cMono(2) were significantly expressed in the blood. These data suggest that there is no enrichment of monocyte subpopulations in blood or liver, consistent with their capacity to patrol the tissues without committing to specific lineages until environmental queues require it^72^.

## Discussion

Advances in technology are shrinking the gap between current analytical capabilities in animal models and humans. Single cell transcriptomics now satisfy the need for characterizing single cell states comprehensively in human samples and has been used to successfully generate single cell tissue maps and characterize disease states. The logical progression of single cell transcriptomics is implementation into relevant clinical trials to investigate the therapeutic effects of novel treatments in a tissue-specific manner. However, the leap to clinical trials presents many practical obstacles around technology infrastructure and variability in data generation. To address these obstacles, we developed a workflow that utilizes the minimally invasive FNA approach to generate data from intrahepatic immune cells at a centralized site without the need for local access to scRNAseq technology platforms.

The FNA approach revolutionizes our access to liver tissue, allowing for frequent, scheduled tissue sampling to collect the most informative time points in disease progression or treatment. The OD450 measurement developed as a quantitative, sample sparing, approach to assess peripheral blood contamination reserves >95% of the FNA sample for scientific investigation. Some practical aspects alone, like assessing the degree of blood contamination, have potential use in current indications for FNAs, such as the cytological evaluation of tumor masses and cysts within organs (thyroid, breast, kidney, lymph node)^73–78^. The full post-sampling workflow could be applied to standard blood samples or to expand the use of FNAs in other tissues, such as kidney, where analysis of early inflammation in transplant recipients through longitudinal sampling could replace serial biopsies^79^.

Our technology comparison verified that using a picowell approach, such as Seq-Well S^3^, effectively captured immune cells without the need for sophisticated equipment. Cellular diversity between the emulsion-based and picowell-based approaches were similar for lymphocytes (T cells, NK cells, B cells) and myeloid cells (monocytes, macrophages) but diverged significantly in their ability to capture granulocytes. The Seq-Well S^3^ picowell technology effectively captured neutrophils in both the blood and liver whereas neutrophils were absent from our microfluidic-based 10x 3’v2 dataset. Human neutrophil capture is a known issue with the 10x 3’v2 data because of their low RNA and high nuclease content, impairing detection within the droplets. Newer versions of the 10x Genomics reagents may improve on neutrophil capture but our data suggest that picowell strategies were more effective. As a result, we identified liver-specific differential transcriptional profiles in neutrophil subtypes involving cytokines and lectins known to modulate the adaptive response in other settings, where they were essential for the activation of an IFN-γ-dependent tumor resistance pathway through the polarization of CD4^-^ CD8^-^ unconventional αβ T cells^80^. This presents a significant opportunity to investigate the role of neutrophils in HBV infection, particularly in liver damage, where the only existing data was generated in mouse models that do not support HBV infection^28, 29, 81^.

An additional benefit of using a picowell-based technology was the ability to load fresh cells onto the arrays locally, freeze the loaded arrays and ship them. This allowed us to use a central laboratory for whole transcriptome amplification, library generation and sequencing at a later time point. We tested the stability of this process on multiple donors by loading and processing parallel arrays fresh, onsite, or in the central laboratory. In our multi-site international collaboration, we found minimal impact on the technical metrics of transcripts, genes and cell capture. We found a maximum of seven differentially expressed genes between matched fresh and frozen arrays. These data demonstrate that we did not introduce technical artifacts in the freezing process and retained the important biological data, including neutrophil capture. This transforms the ability to deploy scRNAseq approaches in clinical studies where tissue is accessible through expanding use of FNA sampling but limited on-site technological infrastructure might otherwise prevent this approach.

Clinically reliable biomarkers in the blood must be the ultimate goal for diseases affecting hundreds of millions of people, such as chronic HBV infection. The lack of immunological biomarkers for disease progression and functional cure has severely hampered therapeutic progress for chronic hepatitis B, which are likely to include immune-based therapies. We deployed the optimized workflow within our collaboration to demonstrate the ability to perform a comprehensive comparison of the liver and peripheral blood immune system across multiple international sites. As hypothesized, we observed tissue compartment specific effects, both in terms of cell enrichment and transcriptional profiles within individual clusters. Of all the lymphocyte populations analyzed, CD8 T cells, known mediators of viral clearance and non-specific liver damage, displayed the most unique transcriptional profiles between the blood and liver. In the five distinct CD8 T cell clusters that we identified, GZMK CD8 T cells displayed the chemokine receptors CCR5 and CXCR6, which are associated with homing to the liver^82^. In addition, GZMK CD8 T cells were the only cluster of cells positive for the IFN-γ transcript. This suggests a potential role in antiviral immunity in the HBV infected liver and/or the potential to drive pathogenesis through induction of IFN-γ regulated chemokines CXCL-9 and CXCL-10.

In contrast to CD8 T cells, CD4 T cells, B cells and NK cells did not show similarly pervasive compartment-specific transcriptional profiles. However, it should be stated that our study was not powered to discover all differentially expressed genes. In addition, our analysis used stringent thresholds for identifying compartment specific gene expression. Genes specifically expressed in the blood or liver were compared while taking the participants into account as covariates. This was necessary to avoid bias towards individual participants but resulted in lower statistical power to detect differentially expressed genes between blood and liver samples, especially in this relatively small cohort. Given the heterogeneity of patients enrolled in our study, we anticipate lower inter-patient variability when patients are selected by defined inclusion/exclusion criteria and more robust transcriptional changes when comparing longitudinal samples from individual patients.

Liver macrophages have been characterized in studies from healthy livers^19^, livers with cholestatic liver disease^83^, and cirrhosis^20^, all of which required tissue digestion. The FNA sampling approach captured macrophages without the need for tissue digestion, minimizing processing time outside of the liver to preserve their in vivo transcriptional profiles. However, macrophage capture was not systematic and varied between patients using the FNA sampling approach. Therefore, calculations such as the frequency of populations would be prone to error, but their transcriptional profiles can be highly informative of the inflammatory environment. In a heterogeneous disease such as chronic hepatitis B, we expected to find a heterogeneous macrophage population, and did so, identifying five different clusters, all of which could be identified from previous reports. Some macrophage populations displayed markers of monocyte to macrophage differentiation, suggesting datasets such as these could be used to investigate the transition of liver infiltrating monocytes. However, because of variability in collection, macrophage-related data should be validated in tissues sections, such as core biopsies, where their transcriptional profiles can be validated by spatial transcriptomics and their frequency and localization can be assessed by fluorescent microscopy. Given their key role in regulating the inflammatory tissue environment, the ability to capture macrophages in FNAs has the potential to provide important insight into the immunological status of the liver.

We found the greatest diversity of cellular phenotypes within short-lived monocytes and neutrophils. Monocytes circulate for approximately a week in the blood, neutrophils even shorter^84, 85^. Because of the short-lived nature of monocytes and neutrophils, they are dynamically regulated within the tissues and periphery by changing environmental cues. Therefore, these cell types may be ideal sentinels for immunomodulatory therapies that induce inflammatory or polarizing cytokines. The FNA procedure, properly timed after treatment, could be used to interrogate the intrahepatic response to immunomodulation to identify more robust biomarkers related to antiviral immunity and immune activation.

In addition to assessing the impact of therapeutic interventions in clinical trials, the ability to electively sample the liver for scRNAseq analysis will open the opportunity to re-define the classical stages of chronic hepatitis B. Historically, chronic hepatitis B has been defined based on the viral load, the presence or absence of HBV antigens (HBeAg) and liver damage (ALT)^86^. While such staging has been relevant for the clinical management of chronic HBV patients, it may obscure effective use of novel therapies. The immune system is anticipated to play a key role in functional cure for new therapies that reduce HBV antigen production (siRNA, antisense oligonucleotides, secretion inhibitors, transcription inhibitors) or that directly target immune cells (vaccines, Toll-like receptors agonists, checkpoint inhibitors). In this case, knowing the status of HBV antigens in the circulation (clinical biomarkers) does not provide insight into the immunological response associated with their change. The workflow we present here opens the possibility to provide single-cell resolution of the evolving immunological status of HBV liver disease through transitions between the different levels of viral control. This type of comparison, across clinical disease phases, allows the identification of immune-related cellular and molecular factors associated with HBV control to define the causal role of these associated factors in disease progression and inform the use of new therapeutic interventions.

The use of FNAs to electively sample the liver, and other tissues, will continue to increase in clinical studies. By combining the FNA sample with scRNAseq, we provide a window into tissue-specific immune response at the single cell level, allowing significantly greater insight into immune function or phenotype than what was achievable by conventional analyses such as flow cytometry. Understanding how to interpret the compartment specific data will be a unique challenge for each tissue, disease or treatment because of the specific cell types involved in disease progression. Facilitating the use of elective tissue sampling using the well-established clinical sampling procedure of FNA with state-of-the-art single cell analysis combines power with granularity to answer these challenges. This will create databases across multiple diseases, in different genetic ancestries and populations, that will spawn opportunities for systematic investigation of human health and disease.

## Methods

### Ethical Statement

Peripheral blood and liver FNAs were collected from 35 participants living with chronic hepatitis B at Erasmus MC University Medical Center (Rotterdam, The Netherlands), Toronto General Hospital (Toronto, Canada) and Massachusetts General Hospital (Boston, USA). Three healthy volunteer blood and liver FNAs were collected at the Janssen Clinical Pharmacology Unit (Antwerp, Belgium). All participants provided written informed consent. The study was approved by institutional review boards at all sites.

### Collection of liver fine needle aspirates

Paired liver FNA and blood samples were obtained to compare intrahepatic and peripheral immune profiles. To minimize variation between sites, the liver FNA procedure was carefully standardized between all participating sites and was performed as follows. The position and movement of the liver during respiration was assessed using ultrasound to avoid large blood vessels. The puncture site was cleaned using chlorhexidine and the participant was covered in sterile dressings, leaving only the puncture site exposed. Upon exhalation, a 25G spinal needle (Braun Spinocan) was inserted intercostally into the liver parenchyma. The stylet was carefully removed and a 10 ml syringe (BD Bioscience) was attached to the needle. Liver cells were aspirated from the parenchyma by pulling back the syringe to create negative pressure, while simultaneously advancing the needle approximately 2.5 centimeters into the liver. Finally, the needle was retracted from the participant and moved to a sterile table where approximately 500 µL of cold RPMI 1640 medium without phenol red (Lonza) was aspirated into the syringe. The cell suspension was then transferred to a 5 mL tube (Axygen) and placed on ice immediately. This procedure was repeated 3 times (total of 4 passes), using fresh needles and syringes for each pass.

### Assessment of peripheral blood contamination in FNA passes

After the FNA procedure, 5 µL of each aspirate was lysed with 45 µL of 1x red blood cell (RBC) lysis buffer (BD Bioscience) and the optical density (OD) was measured at 450 nm (OD450) (OD450) as a proxy for the amount of RBC contamination in the FNA. Site specific OD-value thresholds for significant blood contamination were established using OD450 value of FNAs without visible blood contamination and whole blood as a reference. As a reference blood contamination was also evaluated by the relative frequencies of major T cell populations, determined by flow cytometry. Aspirates with the lowest OD value were selected for downstream analysis.

### Red blood cell depletion of fine needle aspirates and peripheral blood

RBCs in the FNA were removed using magnetic bead-based depletion. Briefly, 12.5 µL of EasySep RBC Depletion Reagent (STEMCELL Technologies Inc.) was added to each FNA pass and incubated for 5 min before placing the sample on the magnet for 5 min (EasySep violet magnet, STEMCELL Technologies Inc.). The FNA cell suspension was poured into another 5 mL polypropylene tube and the procedure was repeated. The RBC depleted cell suspension was then manually counted using trypan blue and a haemocytometer (Neubauer Improved, INCYTO).

For paired whole blood was diluted to 1000 µL using MACS buffer (PBS, 1% BSA and 6 mM EDTA), and RBCs were depleted by addition of 25 µL of EasySep RBC Depletion Reagent.) and incubated for 5 min before placing the sample on the magnet for 5 min (EasySep violet magnet, STEMCELL Technologies Inc.). The blood cell suspension was poured into another 5 mL polypropylene tube and the process was repeated. The RBC depleted cell suspension was then manually counted using trypan blue and a haemocytometer (Neubauer Improved, INCYTO). Magnetic bead RBC depletion retained granulocytes that are normally lost via density gradient peripheral blood mononuclear cell (PBMC) preparation.

For PBMCs, approximately 50 mL of blood was collected in 8.5ml acid-citrate-dextrose (ACD) tubes by standard venipuncture at the time of FNA collection. PBMCs were isolated by standard density gradient centrifugation using Ficoll.

### Flow cytometric analysis to quantify peripheral immune cell contamination

After the OD450 of individual FNA passes were collected and RBCs were depleted, we quantified the frequency of naïve CD8 and CD4 T cells and mucosal-associated invariant T cells in individual FNA passes. Each FNA pass was stained with a viability dye and the following antibodies for 30 min at 4 °C to identify the above-mentioned immune cell populations: CD3, CD4, CD8, Valpha7.2, CD161, CCR7 and CD45RA. The frequency of each population was calculated as a percent of total T cells in the FNA (CD3+) and plotted against OD450 values.

### Cryopreservation of PBMC and FNAs

Any remaining FNA cells and PBMCs were cryopreserved in KnockOut serum Replacement (KO serum, Gibco) with 10% DMSO. Briefly, freezing media A (KO serum alone) and freezing media B (KO serum with 20% DMSO) were prepared on the day of collection. After centrifuging the cells for 5 min at 300g, supernatant was removed and cells were resuspended in freezing media A. An equal volume of freezing media B was added drop by drop, with gentle mixing, giving a final DMSO concentration of 10%. The samples were aliquoted into 1.5 mL cryovials and cryopreserved by cooling to -80 °C in ‘’Mr. Frosty’’ freezing containers and then moved for long-term storage in -150 °C freezers or liquid nitrogen.

### Seq-Well S^3^ transcriptomic profiling

The Seq-Well S^3^ protocol was performed as detailed previously, with several adjustments to improve clinical utility^87^. After RBC depletion, cells were diluted to a concentration of 75,000 cells/mL when possible. A suspension of 200 µL was then loaded onto Seq-Well S^3^ arrays pre-loaded with mRNA capture beads, by adding them dropwise in a zig-zag pattern. When the starting cell suspension was already more dilute, cells were not centrifuged to avoid loss or damage. Instead, an appropriate amount of volume was added in the same dropwise fashion to achieve the same total cell number. This resulted in a larger volume of cell suspension on top of the array, and extra care was taken to make sure that the solution remained on top of the array.

Following membrane sealing for 30 min at 37 °C, samples underwent one of two possible paths. Fresh samples were directly processed on site through cell lysis, hybridization, bead isolation, and reverse transcription as previously described^87^. Frozen samples were placed horizontally in 50 mL conical tubes and immediately stored at -80 °C. Up to two weeks later, the ‘frozen’ samples were placed into complete lysis buffer, the top slides were removed, and the samples proceeded with hybridization, bead isolation, and reverse transcription as previously described. Reverse transcription was performed the same for all samples: 30 minutes at room temperature followed by overnight (18 hr) at 52 °C, both with end-over-end rotation (Hula Mixer, Thermo fisher). After reverse transcription, samples underwent exonuclease treatment, second strand synthesis, whole transcriptome amplification, and Illumina Nextera XT Library preparation. Two SPRI bead-based PCR clean-up steps were performed following both the WTA and Illumina Nextera XT Library Prep Kits library preparation steps. Each time, first a 0.6x and then a 0.8x volume ratio cDNA:SPRI-bead was performed. Sequencing was performed on either a NextSeq 500/550 instrument with a High Output Flowcell and a 75-cycle kit (PE 20/50) or a NovaSeq 6000 instrument with a S4S4 Flowcell and a 100-cycle kit. Samples were demultiplexed according to the Illumina protocols and indices used. Samples were sequenced to an average depth of 1M reads per Seq-Well array.

### 10x genomics transcriptomic profiling

Samples were prepared as outlined by 10x Genomics Single Cell 3’ Reagent Kits v2 user guide. Briefly, samples were washed two times in PBS (Life Technologies) + 0.04% BSA (Miltenyi) and re-suspended in PBS + 0.04% BSA before sample viability was assessed using a haemocytometer (Thermo Fisher Scientific). Following counting, the appropriate volume for each sample was calculated for a target capture of 2000 or 3000 cells. Samples that were too low in cell concentration as defined by the user guide were washed, re-suspended in a reduced volume, and counted again using a haemocytometer prior to loading onto the 10x single cell A chip. After droplet generation, samples were transferred onto a pre-chilled 96-well plate (Eppendorf), heat sealed and incubated overnight in a Veriti 96-well thermocycler (Thermo Fisher). The next day, sample cDNA was recovered using Recovery Agent provided by 10x Genomics and subsequently cleaned up using a Silane DynaBead (Thermo Fisher Scientific) mix as outlined by the user guide. Purified cDNA was amplified for 12 cycles before being cleaned up using SPRIselect beads (Beckman). Samples were diluted 4:1 (elution buffer (Qiagen) : cDNA) and run on a Bioanalyzer (Agilent Technologies) to determine the cDNA concentration. cDNA libraries were prepared as outlined by the Single Cell 3’ Reagent Kits v2 user guide with modifications to the PCR cycles based on the calculated cDNA concentration. The molarity of each library was calculated based on library size as measured with the Bioanalyzer and qPCR amplification data (Sigma). Samples were pooled and normalized to 10 nM, then diluted to 2 nM using elution buffer (Qiagen) with 0.1% Tween20 (Sigma). Each 2 nM pool was denatured using 0.1N NaOH at equal volumes for 5 minutes at room temperature. Library pools were further diluted to 20 pM using HT-1 (Illumina) before being diluted to a final loading concentration of 14 pM. 150 µl from the 14 pM pool was loaded into each well of an 8-well strip tube and loaded onto a cBot (Illumina) for cluster generation. Samples were sequenced on a HiSeq 2500 with the following run parameters: Read 1 – 26 cycles, read 2 – 98 cycles, index 1 – 8 cycles.

### Single-cell RNAseq data preprocessing

Seq-Well S^3^ libraries were pre-processed using the DropSeq pipeline with the tools v2.3.0 as previously described^87^. Briefly, pooled libraries were demultiplexed using bcl2fastq v2.20.0.422 with the following settings: *mask_short_adapter_reads=15* and *minimum_trimmed_read_length=35*. Read alignment was done using STAR 2.5 and the human genome assembly reference GRCh38 (hg38). Aligned cell by gene matrices for each sample were merged across all conditions tested and participants. Preprocessing, alignment, and data filtering was applied equivalently to all samples. Cells with less than 500 UMIs or less than 300 genes were removed from downstream analysis.

Data was log-normalized with a scaling factor of 10,000. The top 2,000 most variable genes as determined by the ‘vst’ method implemented as the FindVariableFeatures function were selected and scaled using a linear model implemented as the ScaleData function, both in the Seurat (v3.1.5) package with version 4.0.2 of R programming language. After, principal component analysis (PCA) was run, the number of significant principal components (PCs) to be used for downstream cell clustering was determined using Jackstraw with a p-value cut-off of 0.05. The best resolution for clustering was determined using an average silhouette scoring across all clusters, testing 40 resolutions between 0.1 and 2 as previously implemented in Ziegler et. al^88^. Marker genes for each cluster were calculated using the FindAllMarkers function (method = ‘wilcox’) implemented in Seurat and each cluster was iteratively subclustered further using the same approach. Subclustering was stopped when the resulting clusters were not meaningfully different. Clusters were annotated as cell type populations based on the expression of genes that are known markers of specific cells. Final marker genes for each intermediary and refined cell type were determined using FindAllMarkers method = “Wilcox” and can be found in Supplementary Table 1.

### Differential frequency analysis between blood and FNA

For each annotated cell population, cell type frequencies per participant were calculated and compared between FNA and blood (whole blood or PBMC, except where noted). Participants with ‘0’ cells within a cell population at the main clustering level were excluded from the analysis. Cell type frequencies were calculated per participant as the fraction of cells within a cell population – e.g., the number of *TCL1A*+ naive B cells within the whole B cell population. The Wilcoxon rank sum test was used to compare the frequency of each cell population across FNA and blood samples and *p*-values were Bonferroni corrected for multiple hypothesis testing.

### Differential expression analysis between blood and FNA

Differential expression analysis was performed to compare the transcriptomic profiles between blood and liver FNA samples within each cell population using the same subset of cells as described for the differential frequency analysis. For each group, a minimum of 5 cells was required to ensure that the samples size was sufficient for the analysis. Read counts were normalized with log2(count+1), and normalized values smaller than ‘1’ were set to ‘0’. The R-package *MAST* was used to obtain hurdle *P*-values which were Bonferroni corrected for multiple hypothesis testing. To remove participant-specific bias in the analysis, participant IDs were treated as covariates. Cohen’s D effect sizes were calculated with R-package *effsize*. Significant genes were determined by corrected *P*-values < 0.05 and Cohen’s D cutoff for each cell type were calculated as mean + 2 x standard deviations of Cohen’s D for all genes.

### Gene set enrichment analysis

Genes that were differentially expressed between blood and liver FNA, cohen’s D > 0.5 were assessed for the overrepresentation of gene sets related to biological states or processes. For this enrichment test, 50 hallmark gene sets with gene symbols were downloaded from the Molecular Signatures Database (MSigDB; http://www.gsea-msigdb.org/gsea/msigdb/genesets.jsp). *P*-values were calculated based on permutations using R-package *fgsea* and multiple hypothesis corrected with the Benjamini-Hochberg method.

## Supplemental Figures

Supplemental Figure 1. Flow cytometry gating strategy to identify naïve CD4 T cells, naïve CD8 T cells and MAIT cells in liver FNA samples.

Supplemental Figure 2. Seq-Well S^3^ vs. 10x Genomics quality control metrics. **(a)** Number of transcripts **(b)** number of genes and **(c)** cell captured from liver FNAs by technology. Number of transcripts from each cell showed a significant difference, paired Student’s T Test for difference of the mean of medians p=0.044. Number of genes from each cell in liver FNAs showed a significant difference, paired Student’s T Test for difference of the mean of medians p=0.013. Number of cells from liver FNA sample showed a significant difference, paired Student’s T Test for difference of the mean p=0.039. **(d)** Number of transcripts **(e)** number of genes and **(f)** cell captured from PBMC by technology. Only the number of genes show significant differences in the blood; paired Student’s T Test for difference of the mean of medians p=0.034. Seq-Well colors orange, 10x colored in blue.

Supplementary Figure 3. CD4 T cell composition in liver vs blood. **(a)** scRNAseq UMAP for CD4 T cells colored by cluster IDs. **(b)** scRNAseq UMAP colored by cluster and split based on tissue of origin, i.e. liver and blood. **(c)** Comparison of cell frequencies between blood and liver within sample (connected through grey lines) for each CD4 cluster. For determining significant differences, Wilcox test with Bonferroni correction was used (adjusted p-value < 0.05). ITGB1+ and CCR7+ CD4 are significantly more in FNA and blood, respectively. **(d)** Dot plot showing top 10 marker genes for each cluster ID. **(e)** Violin plot showing the top 5 marker genes for CD4 T cell clusters. **(f)** Volcano plots depicting differences in gene expression CD4 T cells between compartments. Positive cohen’s D value suggests higher expression in liver. Cohen’s D cutoff calculated as mean + 2 x standard deviation of cohen’s D values of all genes. (H) Hallmark genesets enriched by CD4 T cell clusters. X-axis represents signed log10 of adjusted P-value for the genesets, and positive value suggests enrichment in liver.

Supplementary figure 4. NK cell composition in liver and blood. **(a)** scRNAseq UMAP for NK cells colored by clusters. **(b)** Violin plot showing relevant marker genes for NK cell clusters. **(c)** Dot plot showing top 10 marker genes for each cluster ID. **(d)** scRNAseq UMAP colored by cluster and split based on tissue of origin, i.e., blood and liver. **(e)** Comparison of cell frequencies between blood and liver within sample (connected through grey lines) for each NK cell cluster. For determining significant differences, Wilcoxon rank sum test with Bonferroni correction was used (*Padj* < 0.05). NCAM1-bright EOMES+ NK cells are significantly more present in liver. **(f)** Volcano plots depicting differences in gene expression in NK cells between compartments. Positive Cohen’s D value suggests higher expression in blood. Cohen’s D cutoff is calculated as mean + 2 x standard deviation of Cohen’s D values of all genes. **(g)** Hallmark genesets enriched by NK cell clusters. X-axis represents signed log10 *Padj* for the genesets and the Positive values indicates enrichment in the liver.

Supplementary figure 5. B cell composition in liver and blood. **(a)** scRNAseq UMAP for B cells colored by clusters. **(b)** Violin plot showing relevant marker genes for B cell clusters. **(c)** Dot plot showing top 10 marker genes for each cluster. **(d)** scRNAseq UMAP colored by cluster and split based on tissue of origin. **(e)** Comparison of cell frequencies between blood and liver within sample (connected through grey lines) for each B cell cluster. For determining significant differences, Wilcoxon rank sum test with Bonferroni correction was used (*Padj* < 0.05). All B cell clusters show similar cell frequencies between liver and blood. **(f)** Volcano plots depicting differences in gene expression in B cells between compartments. Positive Cohen’s D value suggests higher expression in blood. Cohen’s D cutoff is calculated as mean + 2 x standard deviation of Cohen’s D values of all genes.

Supplementary Figure 6. Monocyte comparison in liver and blood. **(a)** scRNAseq UMAP for monocytes colored by cluster IDs. **(b)** Dot plot showing top 6 cluster defining genes in each cluster. **(c)** UMAP dimensionality reduction of monocytes by compartment. **(d)** Frequency of monocytes between blood and liver compartments. For determining significant differences, Wilcoxon rank sum test with Bonferroni correction was used (*Padj* < 0.05). Participants with 0 cells within a cell population were excluded. **(e)** Volcano plots depicting differences in gene expression in monocytes between compartments. Positive Cohen’s D value suggests higher expression in blood. Cohen’s D cutoff is calculated as mean + 2 x standard deviation of Cohen’s D values of all genes. **(f)**. Hallmark genesets enriched in monocyte clusters. X-axis represents signed log10 *Padj* for the genesets and the Positive values indicates enrichment in the liver. Classical monocytes; Int. Mono, intermediate monocytes; ncMono, nonclassical monocytes; IFN-stim. cMono, Interferon stimulated classical monocytes.

