## Supplementary Figures for "Clinical implementation of single-cell RNA sequencing using liver fine needle aspirate tissue sampling and centralized processing captures compartment specific immuno-diversity"

#### Suppl Fig 1. FNA Quality Control Gating Strategy

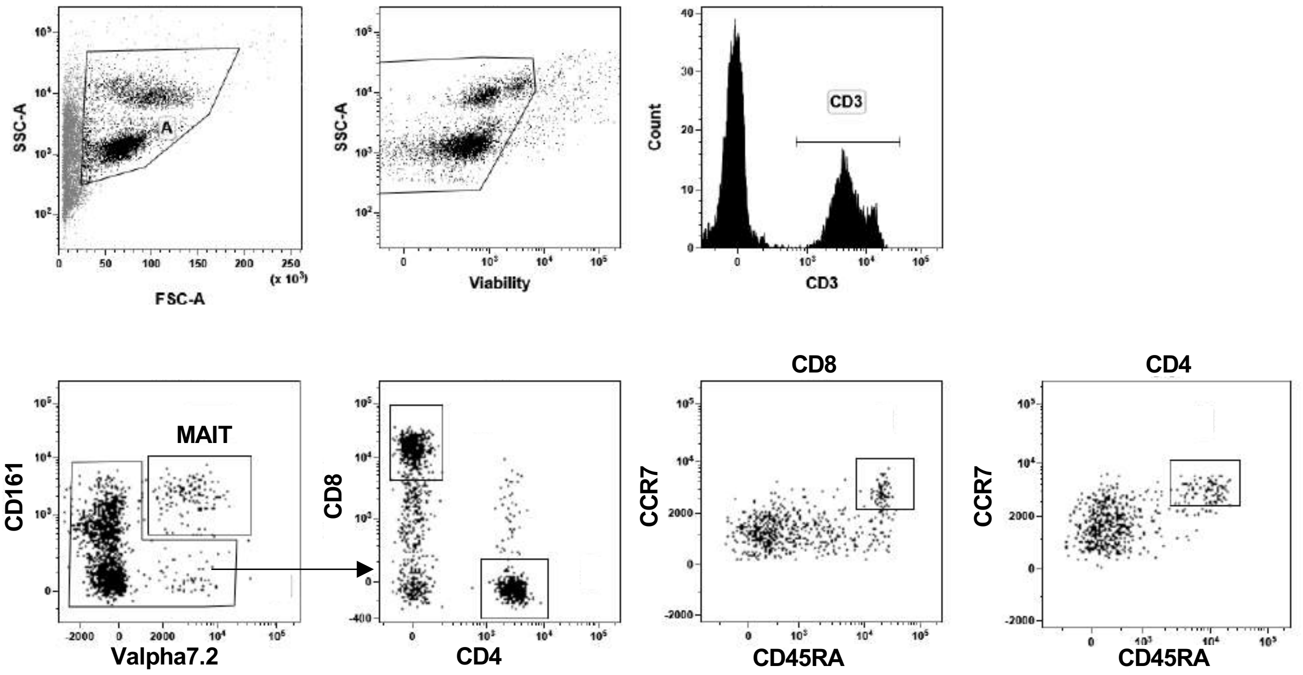

### Suppl Figure 2. scRNAseq QC metrics

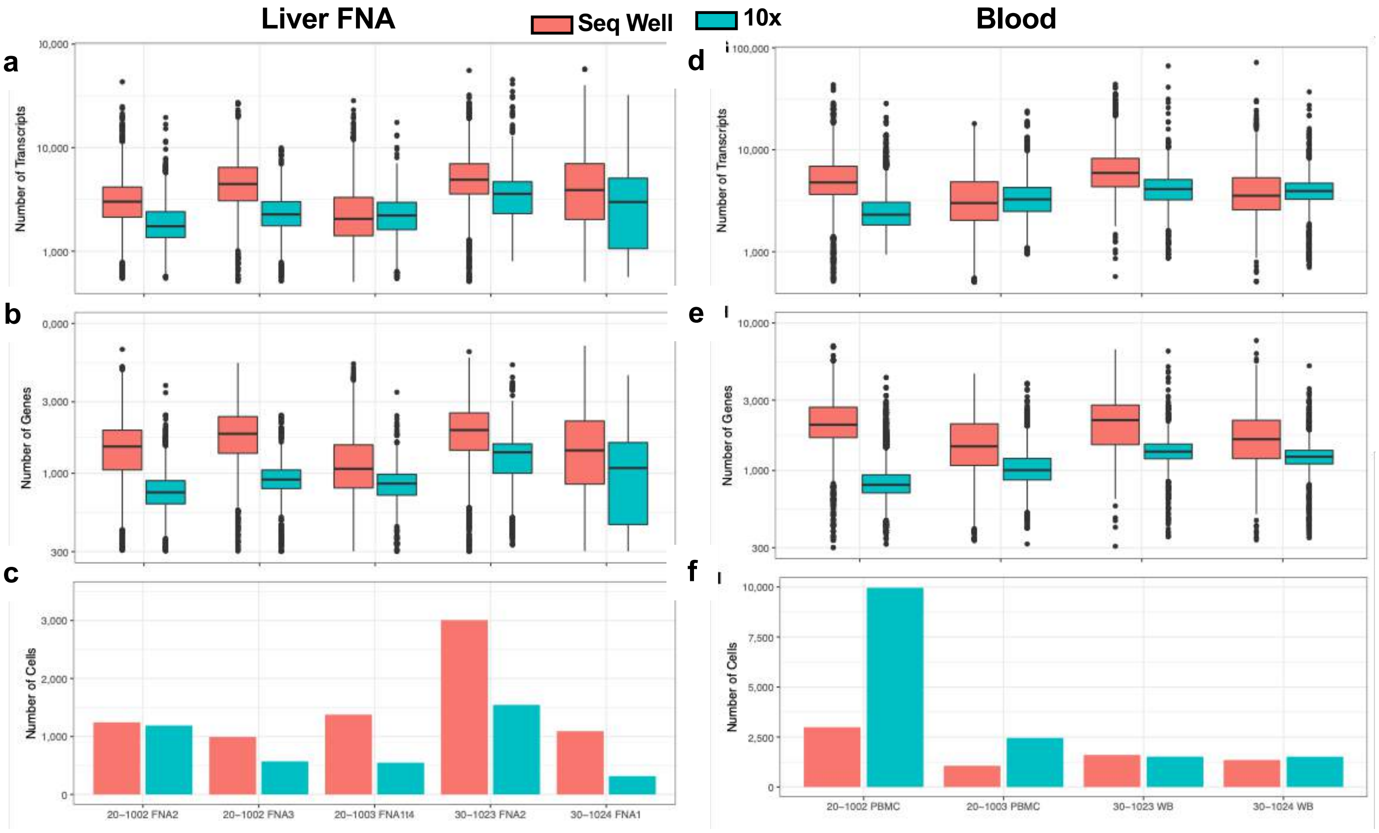

### Suppl Figure 3. CD4+ T cells

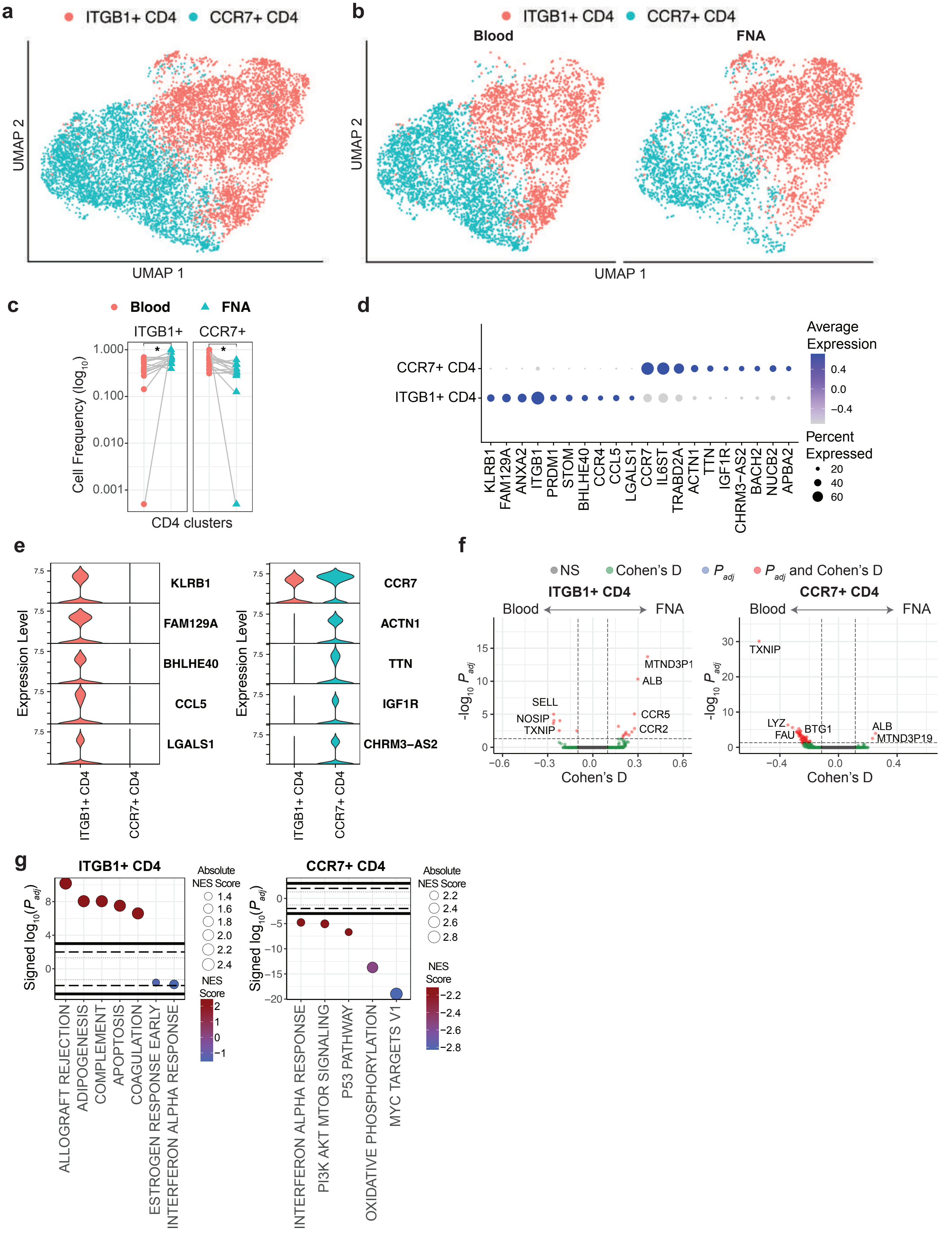

#### Suppl Figure 4. NK cells

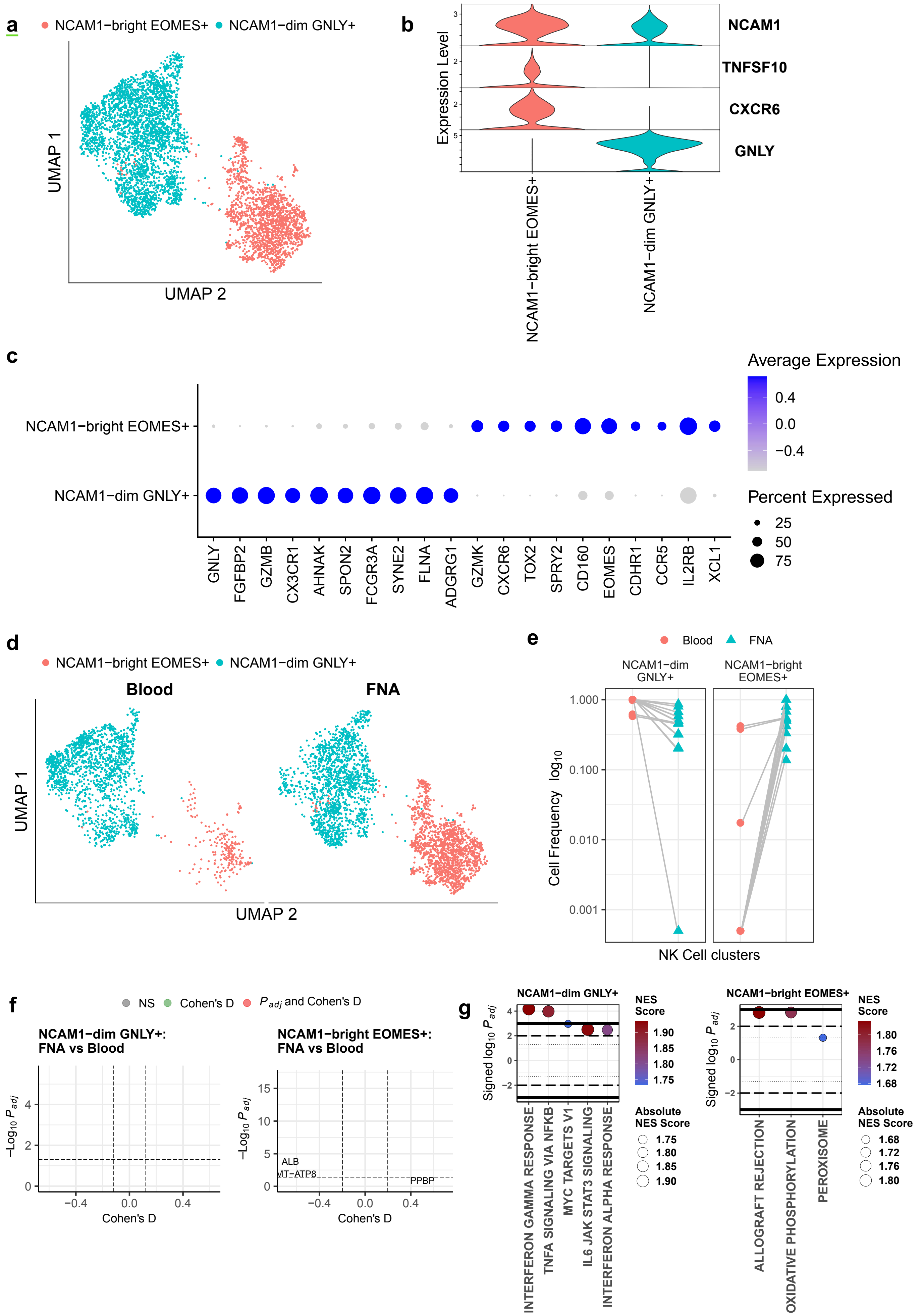

### Suppl Figure 5. B cells

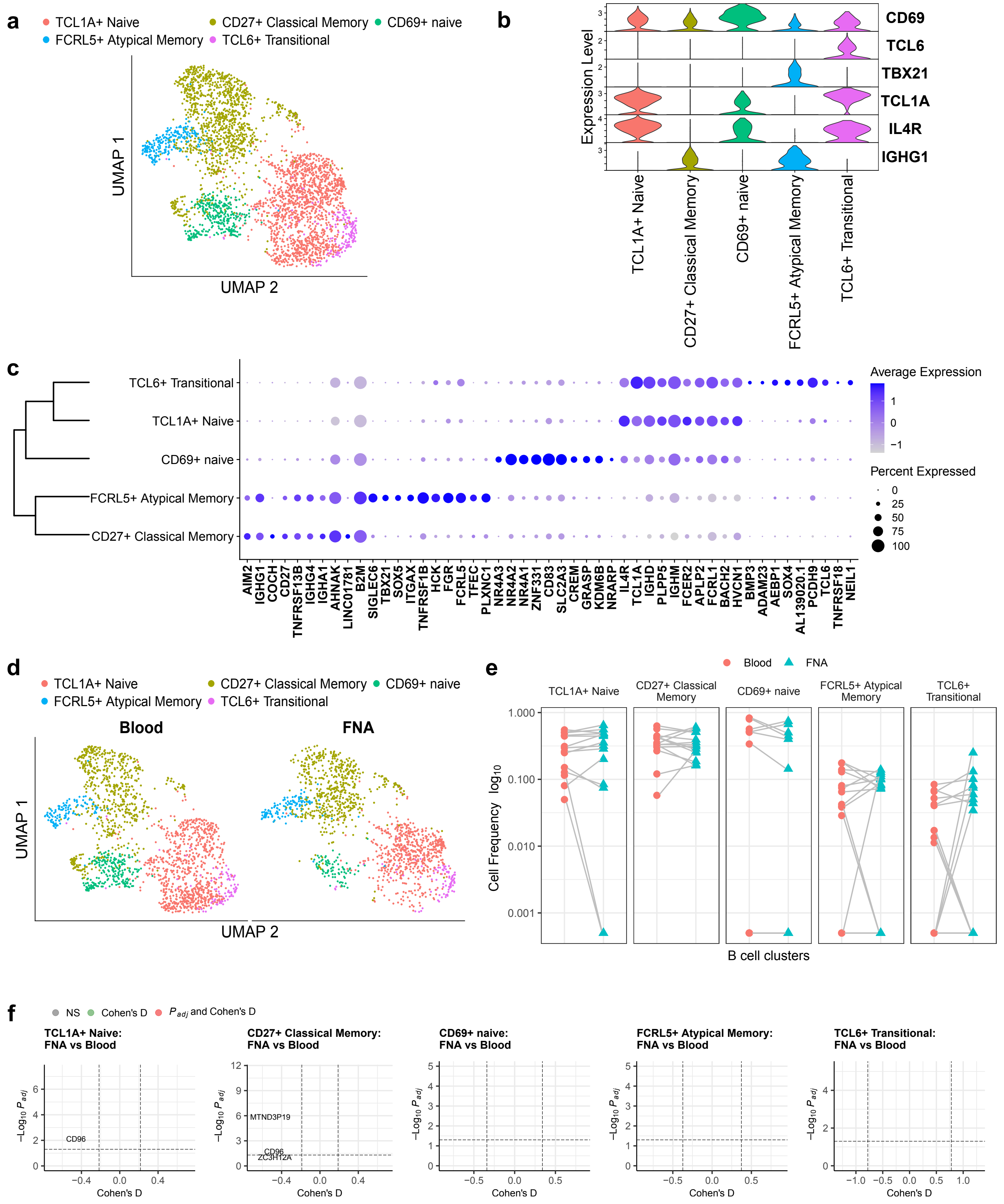

### Suppl Figure 6. Monocytes

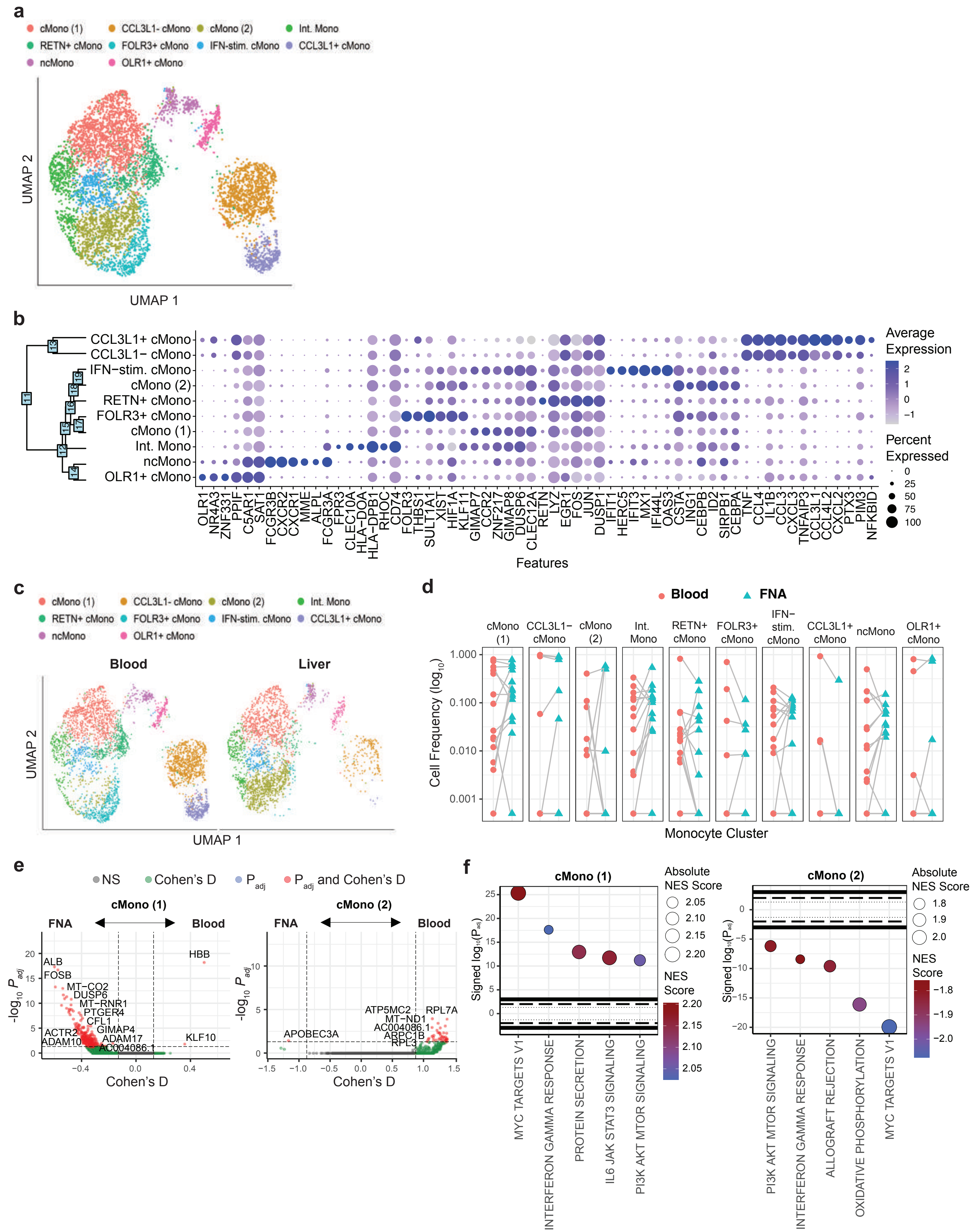
